# Changes in Body Composition, EMG, Kinematic and Kinetic Performance Following Velocity-Based Training vs. 1RM-Percentage-Based Training

**DOI:** 10.64898/2026.01.21.700976

**Authors:** Jairo A. Fernandez Ortega, Laura P. Prieto Mondragon, Hugo M. Borges Sarmento

## Abstract

The aim of this study was to investigate the effects of velocity-based training (VBT) and percentage-based training of one repetition maximum (PBT), on changes in muscle mass (MM), mineral bone density (BMD), and bone mineral content (BMC), maximal squat strength (FSQ), countermovement jump (CMJ), pedaling power (PP), sprint (RV30) and neuromuscular response (EMG). Eighty-five men were randomized to VBT and PBT, and performed full squat (SQ) training three times per week for 12 weeks. Results: RT produced significant increases in FSQ, CMJ, RV30, PP, MM, CMO in both groups, and in BMD only in the VBT group. No significant changes in EMG activity were observed in either group. Significant differences were observed between VBT and PBT for BMD, PP, CMJ, and RV30, with statistical significance set at p<0.05. In conclusion, the VBT group showed better results than the PBT group in the different variables with a lower training load.

## Introduction

Resistance training (RT) has traditionally been based on the percentage of one repetition maximum (1RM) and has been widely used to prescribe and monitor exercise intensity. In recent years, this method has been one of the most accepted tools for prescribing and designing programs due to the ease of programming absolute and relative intensities (RI) into training sessions. However, during the training process, day-to-day and session-to-session variations in 1RM values occur due to fatigue or the rapid increase caused by training adaptation. There is no guarantee that the loads (%1RM) used in each training session represent the planned loads [1]. In general, studies using %1RM approaches cannot provide information on the actual intensity performed during each training session because athletes program IR based on 1RM as a reference to prescribe training [2].

In response to this problem, velocity-based RT (VBT) is a method that can objectively modify the training load variables within a training session to match the previously programmed training [3]. Velocity can accurately determine a %1RM value throughout the load-velocity relationship (r=0.97-0.99) [4,5]. Instantaneous velocity information can be used to: objectively manipulate training load and volume within a session based on the athlete’s performance that day (i.e., VBT training) [6], identify velocity ranges/targets and training loads that can improve training specificity [7,8], identify acute or chronic fatigue or strength gains [5]. A feature of this training is the maintenance of concentric velocity within predetermined ranges (20% maximum decrease) throughout all repetitions and sets.

Only three studies have compared the effects of VBT vs. PBT on kinetic and kinematic variables [1,9,10]. A 6-week training study (2 sessions per week) found that the VBT group showed moderate improvements in maximum strength and jump height compared to the PBT group, which showed only small improvements in maximum strength and nonsignificant increases in jump height [11]. On the other hand, a 7-week RT program (2 sessions per week) was implemented with elite junior rugby athletes and reported similar changes in maximal strength, sprint times, and jump height with CMJ between VBT and PBT [10]. However, one study has some methodological concerns, such as the difference in uncontrolled loads performed during daily field training, on the other hand, most of the training exercises did not use VBT methods and the intervention was not a periodized mesocycle. This study involved twenty-four endurance athletes for 6 weeks, three times a week. The PBT group trained with fixed relative loads that varied between 59% and 85% of 1RM. The VBT group had a target velocity per session prescribed from individualized load-velocity profiles. The results of the study indicated that it was very likely that more favorable effects were observed for VBT than for PBT in CMJ (ES=1.81), 5 m sprint (ES=1.35) and 20 m sprint (ES=1.27); probably favorable in 10 m sprint (ES=1.24) and NDL-COD (ES=0.96) [9].

This is the first study to observe body composition and neuromuscular activation variables, and the population consisted of recreational athletes with no previous strength training experience, indicating an evolution of 1RM during RT, representing a gap between what was programmed and what was performed due to inexperience and possible strength gains due to learning effects. To date, only one study has examined two RT programs, a PBT group with a pre-planned program and another with daily VBT load adjustments in inexperienced subjects [2].

Therefore, the objective of this study was to compare the changes in MM, BMD, BMC, FSQ, CMJ, PP, RV30, and EMG after a training program based on % of 1RM versus another that used VMP to adjust the load of each training session and to monitor the evolution of RI and average velocity achieved during each training session in men with no experience in strength training.

## Methods

### Experimental Approach to the Problem

This was a randomized experimental study with pre- and post-tests administered one week before and one week after the training program. Both groups trained three times a week (48 hours apart) for 12 weeks, for a total of 36 sessions. A progressive RT program consisting only of the FSQ on the Smith machine.

### Subjects

Eighty-five male volunteers (age 20.4 ± 2.94 years; body mass 65.3 ± 8.01 kg; height 171.0 ± 6.9 cm) participated in this study. The recruitment process for the study began in February 2024 and ended in April 2024. Their initial 1RM strength for the full (deep) squat (SQ) exercise was 70.7±13.0 kg. All participants were physically active college students with no experience in strength training. After three weeks of learning the technique, participants had to demonstrate that they could perform a strict and correct technique in order to be included in the current study. Participants were randomly assigned to one of two groups: PBT (n=42) or VBT (n=43). A concealment and double-blind method was used to avoid knowledge of the assignments and to ensure that the evaluators and participants did not know which group they belonged to. Participants had to meet the following criteria: no physical limitations, health problems, or musculoskeletal injuries. None of the subjects were taking any drugs, medications, or nutritional supplements. Two participants in the VBT group and three in the PBT group did not complete 100% of the exercise program and were therefore excluded from the analysis.

#### Ethical considerations

The study was designed in accordance with the deontological standards of the Declaration of Helsinki and approved by the Research Ethics Committee of the University of Applied and Environmental Sciences in session 15. All participants gave their written consent by signing the informed consent document to participate in the study. The study includes minor participants, for which consent was obtained from their parents or guardians. All participants were informed of the benefits and risks of the study before signing the informed consent form to participate in the study.

### Procedures

#### Test Procedures

Prior to each test, a general warm-up was performed using a 5’ bicycle (60 rpm and 60W) and a 5’ specific warm-up for each test. The same warm-up and load progression was used for each subject in the pre- and post-tests. The participants were motivated by the evaluators in all tests to achieve the best possible result.

##### Isoinertial Strength Assessment

1RM in the SQ exercise was assessed by an incremental load test on the Smith machine (Multipower Sport Fitness, USA) without counterweight mechanism. The initial position consisted of knees and hips extended, feet shoulder-width apart, and the bar placed over the trapezius muscle at the level of the acromion to flexion, resulting in a tibiofemoral angle of 35-40° in the sagittal plane, measured with a goniometer (Nexgen Ergonomics, Point Claire, Quebec, Canada) to obtain a FSQ, and after a pause (1-2 s) and at the command of the evaluator, the participant performed an explosive extension at maximum velocity. An eccentric phase control was performed at a speed of ∼ 0.60 to 0.50 m - s - 1. An LPT (T-Force System; Ergotech, Murcia, Spain) was used to measure velocity. The test began with a load of 30 kg and increased by 10 kg to reach an MVP of less than 0.60 m - s, when they began to increase smaller 3-5 kg, determining the 1RM. Three repetitions were performed at the lowest loads (<50%1RM), two at moderate loads (70%1RM), and only one at higher loads (>80%). If participants were able to complete a single repetition at 180° of extension, this was considered the 1RM. Throughout the test, the participants were verbally motivated to do their best, and they observed the displacement velocities in each repetition through a screen. The recovery time between sets was 3’ for the initial loads and 5’ for the final loads [12].

##### Load-Velocity Squat Profile

Seventy-two hours after obtaining the 1RM, the load-velocity profile was determined at 40%-50%-60%-70%-80% of the 1RM and the best MPV was taken at each load. The participants performed the same warm-up protocols as for the 1RM assessment. The specific warm-up for this test was 2 x 8 repetitions with bar weight (13.5kg).

The subjects were required to perform the eccentric phase at a controlled MPV (_0.50_0.65 m/s) and the concentric phase at the maximum intended velocity. This was achieved using a linear velocity transducer (described in detail later), which recorded the kinematics of each repetition and whose software provided real-time visual and auditory feedback [13]. Strong verbal encouragement was provided to motivate participants to achieve maximal effort.

##### Jump, sprint and velocity Tests

Jump height and running velocity can be used as indicators of explosive performance. Participants warmed up as usual and achieved 1RM. Specific warm-up included 3 sets of 10 s jumping with 5 m recovery between sets.

They then performed 5 maximal CMJ with 20 s of rest between each jump. They started from a standing position and performed a knee flexion movement until they approached a 90° angle and immediately pushed off at full speed, always keeping their hands on their hips and their legs extended during the flight time. Participants were given feedback on each jump. The highest and lowest scores were excluded and the remaining scores were averaged for subsequent analyses. Jump height was assessed using the Opto-jump system (microgate Engineering, Bolzano, Italy).

Lower limb performance was assessed using the Wingate test (WAnT) on a Monark 834E cycloergometer (Monark Exercise AB, Vansbro, Sweden). The specific warm-up consisted of 5’ of pedaling at 60-70 rpm at a resistance of 20% of that calculated for the test, and 5’ of acceleration at the end of each minute. After three minutes of rest, the test was performed. Participants were instructed to pedal as fast as possible for 30 seconds against a resistance defined as the product of body mass (kg) by 0.067 kg-body mass-1, applied when each participant reached 75% of the previously obtained RPMmax. The saddle height was adjusted to the height of the iliac spine by verifying that there was no more than 5 degrees of knee flexion with the leg fully extended.

##### 30m Sprint Test

Two maximal sprints were performed over a distance of 30m on an athletic track with a recovery time of 5’ in between. The time was recorded using an infrared light photocell system model WL34-R240 (Sick® Germany) placed at 0 and 30m. The specific warm-up included accelerations of 10, 15, and 20 m with 3’ recovery in between. After 5 minutes of recovery, the test began. The start was made with the front foot 0.5m from the start line where the first photocell was located. The fastest time was used for further analysis.

##### Body Composition

Body composition was assessed before and after the intervention, after an overnight fast (12 h) and 72 h after the last exercise session. Muscle mass, density, and bone mineral content of the lower limbs were assessed by dual-energy x-ray absorptiometry (DXA), with DXA measurements performed using a GE Lunar DXA whole-body scanner (GE Medical Systems Lunar, Madison, WI) and analyzed using software (Lunar encore version 14. 1; GE Medical Systems Lunar) from nine zones encompassing and subdividing the region between the inferior border of the ischial tuberosities and the interline formed at the junction of the femoral condyles and tibial plates in both legs to determine clinically relevant changes in these components. The analyzer was calibrated at the beginning of each day of testing.

##### Electromyography (EMG)

EMG activity of the vastus lateralis (VL) and vastus medialis (VM) muscles of the dominant leg was recorded before, during, and after the RT program in all participants. The EMG was performed with surface electromyography using bipolar Ag / AgCl electrodes of 1 cm diameter (Myotronics noromed, Tukwila, Wash, USA), with a distance between electrodes of 2 cm, placed in the middle of the muscle belly and parallel to the orientation of the muscle fibers, the reference electrode was placed in the proximal bony protuberance, the electrodes were placed on the right leg according to the recommendations established by SENIAM [14]. Skin impedance was reduced to less than 5 kΩ by standard preparation, including shaving, light abrasion, cleaning with alcohol-based gauze, and application of a small amount of conductive gel to each electrode. For VL, the electrodes were placed 2/3 of the way along the anterior superior iliac spine and lateral to the patella. For the VM, the electrodes were placed on the muscle belly at 80% of the distance between the anterior edge of the iliac spine and the anterior edge of the medial ligament. These measurements were performed with the subject seated with the knees slightly flexed and the upper body slightly bent backward; the subject’s thigh with the electrodes in place was photographed to identify the electrode positions and verify the reproducibility of the posttest measurements.

A Motion EMG electromyograph (Mega Electronics Finland) with 6 channels connected via Bluetooth and a range of up to 10 meters was used for recording. The EMG signals were amplified (x100, differential amplifier from 20 to 450 Hz) and digitized at a sampling rate of 2,000 Hz by a personal computer. The raw signals were recorded, full-wave rectified, and filtered (second-order Butterworth low-pass filter with cutoff at 6 Hz: linear envelope). The peak EMG amplitude (mV) and the average EMG amplitude (mV) of the linear envelope were then determined (DATAPAC 2000; RUN Technologies, Laguna Hills, California, USA). For pre- and post-test comparison, the recorded EMG values were standardized to the maximum absolute load lifted. This standardization was performed because absolute EMG values are significantly affected by factors such as subcutaneous tissue thickness, electrode placement (location and orientation), and the method used to shave, scrape, and clean the skin surface.

#### Resistance Training Program

The training program was performed according to the protocol of the study, where the training load was fixed during each session for the PBT group, but adjustable between sets within each session for the VBT group. Warm-up, rest periods between sets (3”), number of repetitions within a set, and number of sets were identical for both groups [9].

The PBT group trained with loads that decreased from session to session during the week (heaviest to lightest) but increased from week to week. For the VBT group, target velocities increased from session to session during the week and decreased from week to week. The researchers supervised all training sessions and recorded and monitored each participant’s adherence to the training protocols. Velocity was controlled with LPT in all sessions for both groups. All participants were required to refrain from any type of strenuous physical activity or training while participating in the research. Participants performed the same warm-up protocols as for the 1RM assessment. Both warm-up and training repetitions were performed at maximum concentric velocity for participants in both groups. For the warm-up, four sets of three squat repetitions were performed using a load of 20% of 1RM. The RT program was the same for both groups, four sets of seven repetitions with three minutes of recovery between sets. All repetitions were monitored and the MPV achieved was recorded.

##### VBT Group Training Program

The VBT group trained with loads adapted to the RI programmed for each session, ensuring the planned goal for the session. In each session, the load to be mobilized was adjusted to obtain the target training velocity for each subject during the concentric phase. To determine the training load of the first set for each VBT session, the MV of the last set of the warm-up (one repetition performed at the training load of the assigned session) was compared with the target velocity. For the other sets, the load adjustment was made with the average MPV of the repetitions of the previous set. In all cases, if the average MPV was 0.06 m s-1 greater or less than the target velocity, the load of the first set was adjusted to ±5%1RM, if it was 0.12 m s-1, a load adjustment of ±10%1RM was made, and so on [15]. Target velocities increased from session to session during the week and decreased from week to week [9].

The velocity of each repetition was controlled by LPT as described above. The session volume consisted of four series of seven repetitions, allowing a maximum loss of 20% of the VMP established for each subject. Once the seven-repetition series were completed, the load for the next series was adjusted according to the average VMP obtained in the series.

##### PBT Group Training Program

The PBT group performed the training with the absolute load of 60-80%1RM (based on their initial 1RM) in such a way that it coincided with the RI programmed for each session, velocity was monitored throughout the training with LPT but was not used for training programming. Loads decreased from session to session during the week and increased from week to week, except for the last four weeks of training when they remained the same [9]. At the beginning of each week, during the first training session, maximal strength was assessed and adjusted if necessary to maintain the programmed IR.

### Statistical Analyses

Analyses of normality of distribution of variables in pre-tests were performed using Shapiro-Wilk tests and homogeneity of variance between groups (VBT vs. PBT) using Levene’s test. Values for all variables are presented as means and ± SD. Absolute test-retest reliability was assessed by CV, while relative reliability was calculated by ICC with 95% confidence interval (CI) using the one-way random effects model method. Statistical significance was set at the P ≤ 0.05 level. Paired-samples t-tests were performed to examine the within-group differences before and after training. In addition, effect sizes (ES) were calculated according to Cohen’s scale, ranging from 0.2 to 0.49 for small effect size, 0.5 to 0.79 for moderate effect size, and 0.8 or greater for large effect size. Statistical analyses were performed using IBMSPSS version 26 software (Chicago, IL, USA).

## Results

At pre-intervention, no significant differences were found between the VBT and PBT groups in the variables analyzed. Descriptive characteristics of the RT program are presented in Table 1.

**Table 1.** Characteristics of the VBT and PBT training program.

| Training Variable | Grupo | Session 1 | Session 2 | Session 3 | Session 4 | Session 5 | Session 6 | Session 7 | Session 8 | Session 9 | Session 10 | Session 11 | Session 12 |
| --- | --- | --- | --- | --- | --- | --- | --- | --- | --- | --- | --- | --- | --- |
| Velocity (m·s<sup>-1</sup>) | VBT | 0,66±0,06 | 0,68±0,03 | 0,71±0,05 | 0,68±0,04 | 0,72±0,03 | 0,74±0,04 | 0,70±0,06 | 0,74±0,06 | 0,78±0,04 | 0,72±0,07 | 0,76±0,03 | 0,80±0,02 |
| %1RM | PBT | ~60%RM | ~56%RM | ~52%RM | ~65%RM | ~61%RM | ~57%RM | ~70%RM | ~66%RM | ~63%RM | ~70%RM | ~66%RM | ~62%RM |
|  |  | Session 13 | Session 14 | Session 15 | Session 16 | Session 17 | Session 18 | Session 19 | Session 20 | Session 21 | Session 22 | Session 23 | Session 24 |
| Velocity (m·s<sup>-1</sup>) | VBT | 0,74±0,03 | 0,78±0,02 | 0,82±0,01 | 0,76±0,02 | 0,80±0,03 | 0,83±0,03 | 0,78±0,02 | 0,82±0,02 | 0,83±0,01 | 0,80±0,02 | 0,82±0,03 | 0,83±0,02 |
| %1RM | PBT | ~70%RM | ~66%RM | ~62%RM | ~75%RM | ~71%RM | ~67%RM | ~75%RM | ~71%RM | ~67%RM | ~75%RM | ~71%RM | ~67%RM |
|  |  | Session 25 | Session 26 | Session 27 | Session 28 | Session 29 | Session 30 | Session 31 | Session 32 | Session 33 | Session 34 | Session 35 | Session 36 |
| Velocity (m·s<sup>-1</sup>) | VBT | 0,82±0,03 | 0,83±0,04 | 0,83±0,02 | 0,78±0,03 | 0,82±0,02 | 0,83±0,03 | 0,80±0,03 | 0,82±0,04 | 0,83±0,01 | 0,82±0,03 | 0,83±0,05 | 0,83±0,03 |
| %1RM | PBT | ~80%RM | ~76%RM | ~72%RM | ~80%RM | ~76%RM | ~72%RM | ~80%RM | ~76%RM | ~72%RM | ~80%RM | ~76%RM | ~72%RM |

Significant increases (p<0.01) were observed after twelve weeks of training in both groups for all study variables, but the magnitude of the improvements was not similar: in the FMSQ (VBT 28.7%; ES=1.71 PBT 29.8%;ES=2.45), CMJ (VBT 15.5%; ES=3.09; PBT 4.2%;ES=1.22), RV30 (VBT 14.1%;ES=1.12; PBT 3.1%;ES=1.0) PP (VBT 30.8%; ES=0.86; PBT 13.8%;ES=0.21) energy production/lean mass (VBT 27.4%;ES=0.78; PBT 9.8%;ES=0.01) MM (VBT 2.89%:ES=1.06; PBT 3.32 %;ES= 0.85)) BMC (VBT 0.69%;ES=0.61; PBT 0.47%;ES=0.32) BMD (VBT 0.70%;ES=0.92) Table 2-4. When comparing the effects of training between the groups Table 4, significant differences (p<0.01) were identified in the changes produced by training, being more beneficial for VBT in: BMD, CMJ, RV30, PP compared to PBT.

**Table 2.**
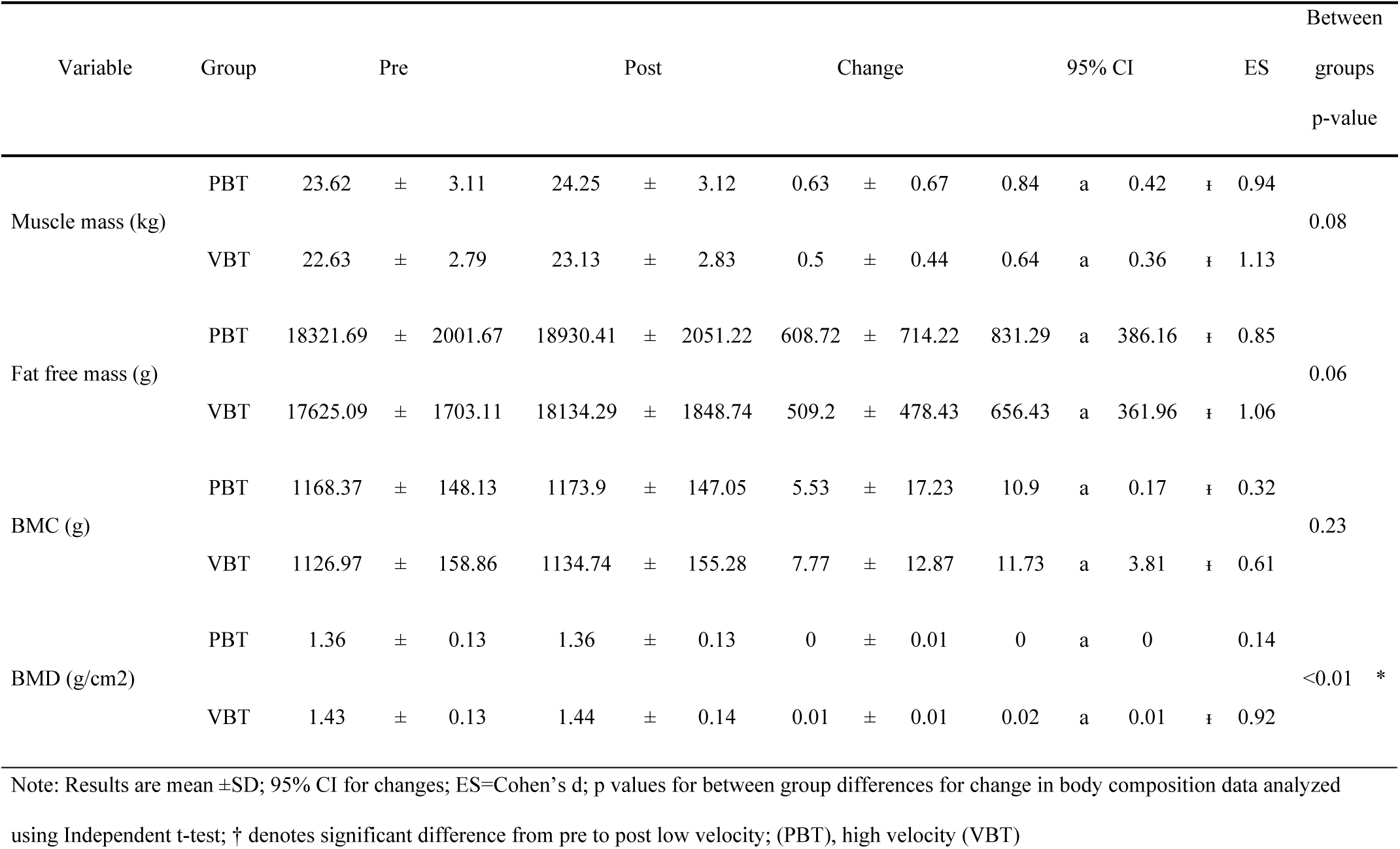
Pre-, post-, mean change, and effect sizes for body composition data.

**Table 3.**
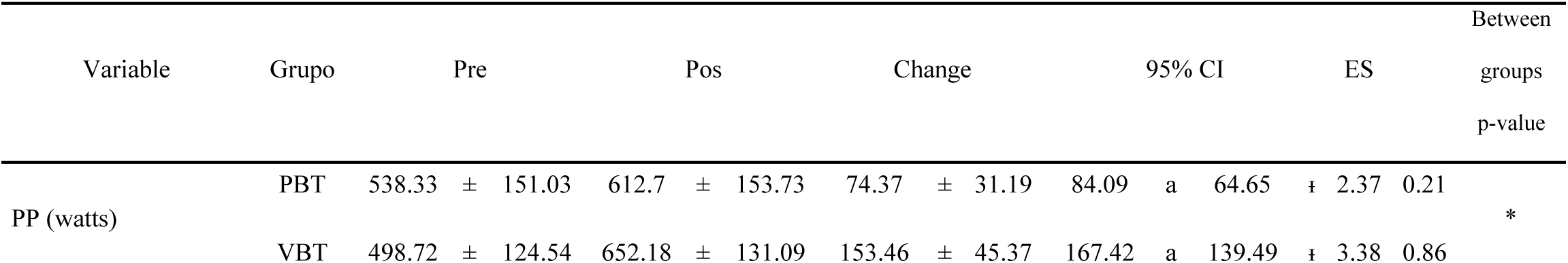
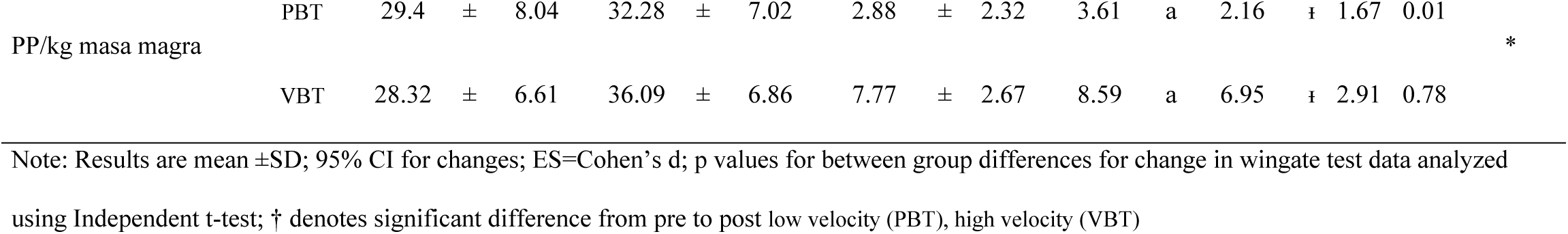
Pre-, post-, mean change, and effect sizes for Wingate testing data.

**Table 4.**
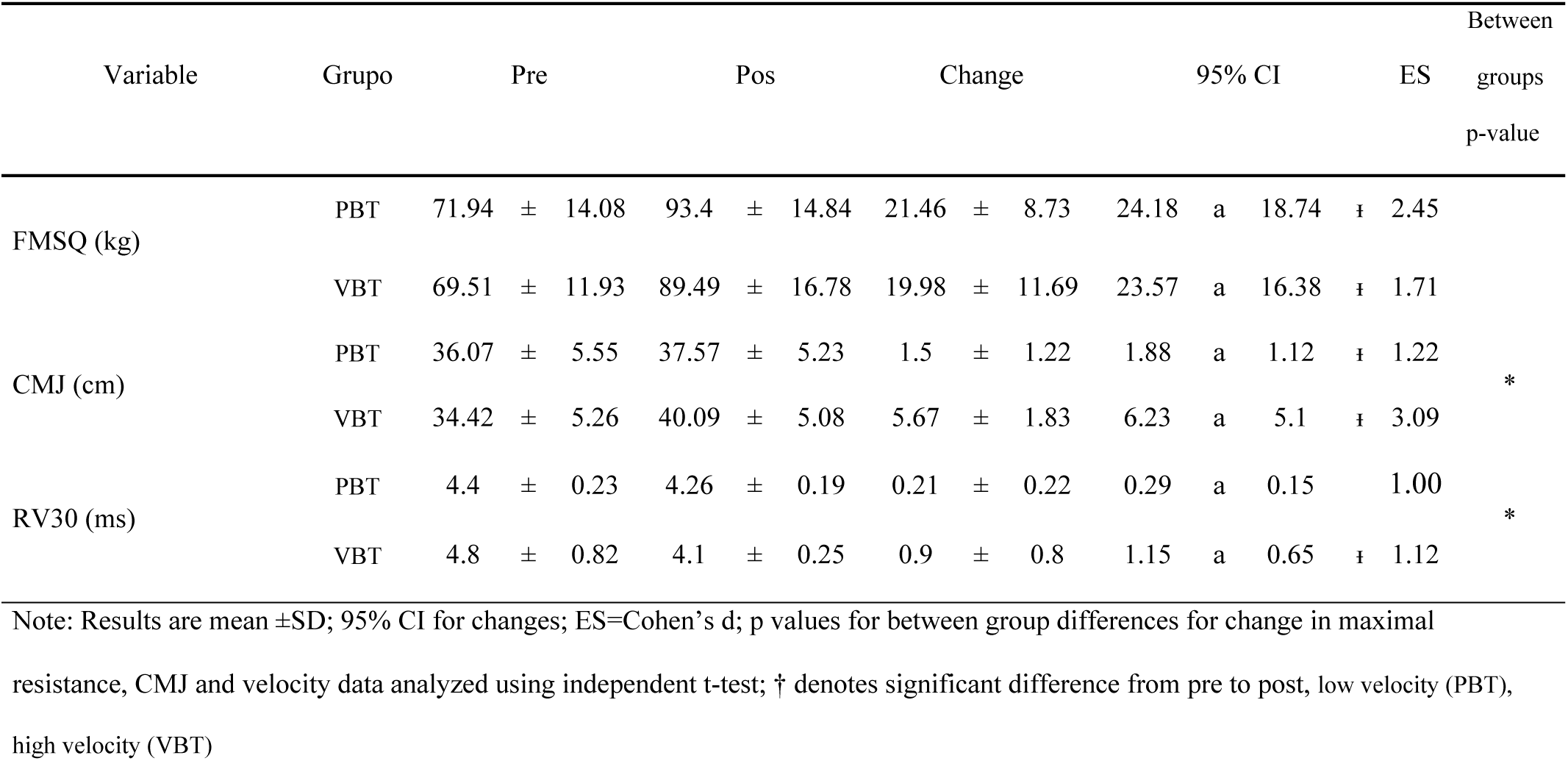
Pre-, post-, mean change, and effect sizes for maximum resistance, CMJ and velocity data.

The increase in the FMSQ did not show significant differences between the two groups after twelve weeks of training. Regarding neuromuscular activity, no significant differences in EMG VL and VM amplitude were observed between the two training protocols. An increasing trend in activation between pre and post was observed in the amplitude of EMG, VL and VM, but the differences were not statistically significant Table 5.

**Table 5.** Maximum EMG (mV) activation values of the VL and VM before and after 12 weeks of exercise.

| Muscle | Group | Pre | Pos |
| --- | --- | --- | --- |
| VL | PBT (n=42) | 1,78 ± 1 | 2,31 ± 1,06 |
|  | VBT (n=43) | 1,75 ± 1,14 | 2,52 ± 1,26 |
| VM | PBT (n=42) | 1,96 ± 1,21 | 2,81 ± 1,17 |
|  | VBT (n=43) | 1,86 ± 1,08 | 2,92 ± 0,98 |

## Discussion

The purpose of this study was to compare two strength training protocols and determine if they would produce different strength and performance outcomes. Several hypotheses were posed, the first being that the VBT group would produce a greater degree of adaptation for CMJ, RV30, PP and EMG compared to PBT due to the individualized approach to the load-velocity relationship of VBT. The results of the present study partially support this hypothesis, and significantly more favorable training effects were observed for the VBT group, but not for EMG. The second hypothesis indicated that PBT training would produce a greater magnitude of adaptation for FMSQ, MM, BMC and BMD, the results reject this hypothesis because no significant favorable differences were observed for any of the groups in FMSQ, MM, BMC, on the contrary, significant differences were more favorable for the VBT group in BMD. Faster average training repetitions may explain the favorable effects of VBT. The main finding of this study was that the VBT group, which performed the same volume (number of repetitions in each training session) as the PBT group but with a lower training load in each session, showed greater gains in running speed, jump height, pedaling power and bone mineral density than the PBT group.

Several studies have reported effects similar to those in the present study [2,16,17]. The stimulus produced by the VBT group, characterized by a low degree of fatigue and a high speed of repetitions within the series, may be sufficient to induce strength and power adaptations in the non-athletic population, highlighting the usefulness of VBT in RT. However, an element to take into account for the present study is that the MPV was evaluated in each series to adjust the training loads according to the programmed MPV, the magnitude of the increases in maximum force for both groups are greater than those reported in other studies (VBT: ∼9%, ES = 0.59; PBT: ∼8%, ES = 0.44) [11], (VBT: 11.3%; ES = 0.89; PBT:12.5%; ES = 1.41) [9] this difference may be due to a greater volume of training, 12 weeks (3 sessions/week) for a volume of 28 repetitions per session, 84 per week, for a total of 1008 repetitions for the entire training intervention, and the characteristics of the population in the present study, not being athletes nor having experience in strength training.

In general, studies using 1RM percentage-based approaches cannot provide information on the actual intensity performed during each training session because athletes use a pre-programmed 1RM-based RI as a reference to prescribe training [2]. In the current study the relative improvement in 1RM was quite similar between the groups, but the PBT group trained at slightly higher loads and slower speeds compared to the VBT group. This slightly lower load for the VBT group allowed the planned speed for the session to be maintained and resulted in large between-group differences (ES = 2.17) in the MPV of training repetitions. However, despite faster repetition rates in the VBT group, there were no significant differences in 1RM gain compared to the PBT group. In this sense, a study observed in a group of men who performed RT on the bench press, for three weeks, twice a week, at maximum concentric velocity in each repetition, stopping when there was a drop in repetition velocity below 20% in relation to the fastest repetition of the first set. Both the VBT and PBT groups increased their 1RM by ∼10%, but it was slightly higher in the VBT group, which was attributed to greater recruitment of high firing rate motor units, which may improve the rate of force development leading to increases in 1RM. However, this was not verified in our study where, despite an increase in electrical activity in the VBT group, it was not significantly different from that in the PBT group [18].

Another important aspect of the present study is that the total training volume was the same in both groups, obtaining similar increases in muscle mass, which confirms the results of the research indicating that different RT programs with similar total load produce similar results in muscle hypertrophy and that the adjustment of the training load in each series generates different stimuli from the early stages [2,19]. Furthermore, the authors observed similar increases in muscle volume with the same two types of training used in the present study [20]. The hypertrophy observed in the present study in the two training groups could also be explained by the increase in EMG amplitude observed after 12 weeks of training. This increase could be due to a higher motor unit recruitment and/or activation rate, which has been identified as a central factor in triggering muscle hypertrophy.

Research on velocity as a variable for muscle mass gains has shown that light-load resistance training (RT) performed at high velocity can produce similar effects to traditional heavy-load RT. The authors examined two types of RT-one at 20% maximal velocity and another at 80% 1RM-on fat mass index (FMI), fat-free mass index (FFMI), and lower limb muscle mass (LLMM). They observed significant increases in LLMM (p<0.01 for both groups) and FFMI (p<0.05 for the speed group) [20]. Similarly, a meta-analysis showed that hypertrophic responses were comparable when repetition durations ranged from 0.5 to 8 seconds [21]. However, most of the studies in the analysis lacked control over the duration of the movement phase (eccentric vs. concentric), limiting definitive conclusions. Furthermore, the current evidence remains inconclusive as to whether movement velocity manipulation specifically affects muscle hypertrophy [22]. Consistent with previous research, this study found significant increases in muscle mass of 2.67% for power-based training (PBT) and 2.21% for velocity-based training (VBT) at 12 weeks, with no significant differences between groups. These results suggest that high %1RM RT is not the only method to stimulate muscle protein breakdown and hypertrophy in untrained adolescents.

Favorable improvements in jump height, running speed, pedaling power, and bone mineral density occurred in the VBT group compared to the PBT group. These findings are consistent with previous research indicating that greater training velocity can lead to superior improvements in jumping and sprinting [9,23] suggesting that mobilizing a load as quickly as possible is a critical factor in improving muscle strength and power. Recently, RT performed at maximal voluntary velocity has been shown to be of paramount importance in maximizing strength gains and athletic performance (jumping ability, running speed, pedaling power) [6]. One study reported a significant increase of 5% in CMJ for the high-velocity training group only [11].

Regarding running speed, the results of this study are consistent with the findings of previous studies, including meta-analyses, which report better results in running speed when RT is performed at low loads and high speeds than when high loads and low speeds are used [20,24,25]. High velocity training results in the recruitment of high frequency firing motor units, which together with possible increased motor unit synchronization would produce neural adaptation. RT performed at high speed causes a greater and/or more effective recruitment of fast-twitch muscle fibers, changes in myosin heavy chain isoform composition, and increases in tendon aponeurosis stiffness, all of which seem to induce substantial improvements in the rate of force development and, consequently, in different types of explosive actions such as sprinting, VJ, and PP [13,26]. In the present study, the group that trained with lighter loads had a higher movement velocity (mean 0.738 m.s-1) during each training set, compared to the group with heavier loads (mean 0.423 m.s-1).

Regarding RT based on %1RM and its effect on bone mineral content, several studies reported that both moderate (40-60% 1RM) and high intensity (70-90% 1RM) RT exercise protocols significantly increased BMD [27,28]. In the present study, a 1% increase in BMC and BMD was observed in the VBT group versus 0.87% in the PBT group, which is consistent with these findings in the aforementioned studies [29]. This contrasts with previous studies that reported that only high-intensity RT performed at high loads (70-80%1RM) was effective in increasing BMD, whereas a low-intensity program caused no change. The differences between studies may be due to the diversity in RT methodology, which differs in intensity, frequency, total volume, duration, or all at the same time, elements that have a great impact on the effects of RT [13]. In most studies, including the present one, the increase in BMD and BMC is modest (0.5% to 2.5%). This may be due to an underestimation caused by DEXA because it only measures bone mass, which is part of bone strength, and is likely to miss structural (internal and periosteal) bone changes [30]. Bone strength is determined not only by bone mass, but also by size, shape, structure and material and collagen properties [30] so it is believed that DXA may underestimate the true effects of mechanical loading on bone [31]. In summary, the results of most studies suggest that with adequate intensity and relatively few repetitions, they are sufficient to generate an adaptive skeletal response of osteocytes [21].

In the present study, significant negative correlations (r=-0.62, p=0.005) were observed between individual relative changes in CMJ height and individual relative changes in RV30. Changes in FMQS show strong associations (r=-0.65) with changes in RV30 performance, results observed in previous studies [9]. They also found similar improvements in the FMSQ, 29.8% in the PBT group compared to 28.7% in the VBT group. These results are consistent with previous observations reported in other studies [17,29]. A non-significant increase in EMG values was also observed in both groups, which was associated with significant increases in maximal strength and muscle power, results consistent with those reported in other studies [32]. This increase in EMG in both groups may be due to the high intensity of both types of training. On the other hand, the greater increase in the VBT group may be explained by the greater force generation required to accelerate the bar at high speeds and for the braking phase of the movement, thus requiring greater motor unit recruitment [33]. These changes in motor unit recruitment or motor unit synchronization have also been proposed as a reason for the changes in EMG values, suggesting that it is possible that changes in firing rate may have contributed to the observed increase in force [34].

In conclusion, our main findings were that VBT training may provide a superior stimulus to induce neuromuscular adaptations that produce greater improvements in VJ, speed over 30 m, pedal force, BMD, and similar or even greater increases in maximal squat strength, muscle mass, and BMC than PBT training. Furthermore, although no statistically significant differences in EMG variables were observed for any training group, only VBT showed slight increases in EMG activity. Therefore, the results of the present study seem to indicate that the establishment of high-velocity, low-load training allows for a more effective and efficient training stimulus, as it produces similar or even greater gains in neuromuscular performance at lower training loads that induce a much lower degree of fatigue (mechanical and physiological stress) compared to higher training loads that induce greater velocity losses.

### Practical Applications

Our results suggest the importance of evaluating the daily load adjustment for a planned IR and the degree of match or mismatch between what is planned and what is performed. By comparing the two strategies, it is possible to accurately assess the specific IR that is being performed and determine the exact moment during the training period when there is a difference between the programmed training and the one performed in inexperienced subjects. In addition, the LVP-VBT method allows individuals to lift appropriate loads on a given day to accommodate individual training adaptation rates.

## Acknowledgments

This study had no funding sources and no conflicts of interest. The results of this study do not constitute an endorsement of any product by the authors or the NSCA.

